# Bloodmeal metabarcoding of the argasid tick (*Ornithodoros turicata* Dugès) reveals extensive vector-host associations

**DOI:** 10.1101/2023.08.07.552345

**Authors:** Sujata Balasubramanian, Rachel E. Busselman, Nadia Fernandez-Santos, Andy Grunwald, Nicholas Wolff, Nicholas Hathaway, Andrew Hillhouse, Jeffrey A. Bailey, Pete D. Teel, Francisco C. Ferreira, Sarah A. Hamer, Gabriel L. Hamer

## Abstract

Molecular methods to understand host feeding patterns of arthropod vectors are critical to assess exposure risk to vector-borne disease and unveil complex ecological interactions. We build on our prior work discovering the utility of PCR-Sanger sequencing bloodmeal analysis that work remarkably well for soft ticks (Acari: Argasidae), unlike for hard ticks (Acari: Ixodidae), thanks to their unique physiology that retains vertebrate DNA from prior bloodmeals viable for years. Here, we capitalize on this feature and apply bloodmeal metabarcoding using amplicon deep sequencing to identify multiple host species in individual *Ornithodoros turicata* soft ticks collected from two natural areas in Texas, United States. Of 788 collected *O. turicata*, 394 were evaluated for bloodmeal source via metabarcoding, revealing 27 different vertebrate host species (17 mammals, 5 birds, 1 reptile, and 4 amphibians) fed upon by 274 soft ticks. Information on multiple hosts for individual *O. turicata* was derived from 168 of these (61%). Metabarcoding revealed more mixed vertebrate bloodmeals in *O. turicata* previously processed using Sanger sequencing. These data reveal wide host range of *O. turicata* and demonstrate the value of bloodmeal metabarcoding for understanding the ecology for known and potential tick-borne pathogens circulating among humans, domestic animals and wildlife such as relapsing fever caused by *Borrelia turicatae*. Our results also document, for the first time an off-host soft tick collected to have evidence of prior feeding on wild pig which is a critical observation in the context of the threat of enzootic transmission of African swine fever virus if it were introduced to the US. This research enhances our understanding of vector-host associations and offers a promising perspective for biodiversity monitoring and disease control strategies.

## 1 INTRODUCTION

Bloodmeal analysis of hematophagous arthropods can reveal local vertebrate diversity (Schnell et al., 2012) and identify host species utilized by vectors that transmit viruses, bacteria and parasites that affect humans, domestic animals, and wildlife (Borland & Kading, 2021). Recently, bloodmeal analysis using a marker amplicon and next generation sequencing (henceforth, metabarcoding) has revealed an increase in number and diversity of hosts utilized by mosquitoes (Logue et al., 2016), triatomines (Balasubramanian et al., 2022; Dumonteil et al., 2018; Murillo- Solano et al., 2021; Polonio et al., 2021; San Juan et al., 2023) and sand flies (Abbasi et al., 2019), providing insights from both the perspective of pathogen transmission and also to elucidate the community of hosts in the environment. For instance, by using this approach (Kocher et al., 2017), Kocher et al., (2023) demonstrated that locations with higher indices of anthropogenic modification were associated with lower mammal diversity, and increased relative abundance of bloodmeals derived from reservoir hosts, which in turn favors vectorial pathogen transmission. Therefore, bloodmeal metabarcoding appears to be a powerful tool to unveil vector-host interactions in areas where pathogens circulate among humans, domestic animals and wildlife.

Soft ticks (Acari: Argasidae) are long-lived, hematophagous arthropod vectors, including approximately 200 species, known to vector a variety of pathogenic bacteria and viruses (Mans et al., 2019; Manzano-Román et al., 2012). The genus *Ornithodoros* includes important vectors of medical and veterinary importance globally. In the USA, *O. turicata* is found in Florida (recognized as *O. turicata americanus* by (Beck et al., 1986)) and from the southwestern US from Texas to California including the high plains of Kansas and Colorado, extending into Mexico (Donaldson et al., 2016; Dworkin et al., 2002). Different species of *Ornithodoros* are important vectors and reservoirs for pathogenic bacteria in the spirochete genus *Borrelia* that cause tick-borne relapsing fever (TBRF) in humans and in domestic animals in the Americas (Faccini-Martínez et al., 2022). *Ornithodoros turicata* transmit *Borrelia turicatae*, an agent of TBRF, which is found in Texas and other western states associated with caves or animal burrows (Beeson et al., 2023; Campbell et al., 2019; Dworkin et al., 2008). Multiple *Ornithodoros* ticks, including *O. turicata*, are also competent vectors of African swine fever virus (ASFV). ASFV causes 100% mortality in domestic swine (Manzano-Román et al., 2012), and is currently considered the most impactful disease for the pig production industry (Jori et al., 2023). ASFV is a DNA virus (family Asfiviridae) originated in Africa that has emerged in Europe, Asia, and the Caribbean (Gonzales et al., 2021; Jean-Pierre et al., 2022; Penrith et al., 2013). There is no broadly-distributed vaccine for ASFV and the current method of control relies on biosecurity measures to prevent contact between the non-infected domestic pigs with virus sources (Brake, 2022; USDA, 2022), including *Ornithodoros* ticks, which can transmit the virus for at least one year after infection (Boinas et al., 2011).

*Ornithodoros* ticks feed on a broad diversity of vertebrates including reptiles, birds, livestock, companion animals (e.g., dogs) and humans (Busselman et al., 2021; Kim et al., 2021; Palma et al., 2013). DNA from prior bloodmeals is retained for years in soft ticks (Beck et al., 1986; Francıs, 1938; Kim et al., 2021), and our recent study using *O. turicata* reared under laboratory conditions confirmed chicken (*Gallus gallus*) and domestic pig (*Sus scrofa domesticus*) bloodmeals via Sanger sequencing 1,105 and 622 days, respectively, after feeding took place (Busselman et al., 2021). The habitat of *O. turicata* includes caves and animal burrows with nidicolous species of vertebrates (Adeyeye & Butler, 1989; Francıs, 1938; Manzano-Román et al., 2012). Because soft ticks can live for many years in environments with favorable temperature and humidity conditions and feed multiple times throughout their lifetime (Dworkin et al., 2002), analyzing their bloodmeals may become an efficient way to characterize local vertebrate host communities.

In this study, a metabarcoding approach was applied to identify the diversity of hosts utilized as bloodmeal sources by *O. turicata* collected from two natural areas in Texas, USA between 2019 and 2022. Understanding the host preferences and blood feeding habits of *O. turicata* contributes to informing preventive and control measure. This tool could also help identify soft tick- vertebrate host networks in other regions of the world such as Africa, Europe, and Asia where ASFV is endemic, which was recently highlighted as an important knowledge gap (Jori et al., 2023).

## 2 METHODS

### 2.1 Locations and soft tick sampling

Soft ticks were collected from two natural areas in Texas, USA. One site was Government Canyon State Natural Area (GCSNA), San Antonio, TX (Lat: 29.549316, Lon: -98.764715; Texas Parks and Wildlife State Park Scientific Study Permit No. R3-02-19). Samples from this location were used for a prior study (Busselman et al., 2021). Sampling took place in March 2019 from three caves (Bone Pile, Little Crevice and Mad Crow). As previously described, dry ice baited sticky traps were placed at the opening of the three caves for two consecutive days with one check and reset (removal of ticks, replacement of dry ice and sticky material) at 24 h. The glue board material (Trapper Max Mouse & Insect, Sheffield, UK) remained sticky even with moisture or rain.

The second sampling site was Laguna Atascosa National Wildlife Refuge (LANWR), Los Fresnos, TX (Lat: 26.22901, Lon: -97.34725; US Fish and Wildlife Service Research & Monitoring Special Use Permit No. 2022-LA-001). Sampling at LANWR took place between February and June, 2022. LANWR was established as a National Wildlife Refuge for migrating and wintering birds with a diverse community of vertebrates including bird, mammal and reptile species. The landscape consists of thornscrub, coastal prairies, salt flats, and estuaries and given the absence of caves in this landscape, traps were placed in 11 animal burrows (designated LA-1, LA-2, LA-6, LA-7, LA-8, LA14, LA15, LA-23, LA-24, LA-25, LA-26). The dry-ice baited sticky traps used at LANWR differed from those used in GCSNA and were CO_2_ baited sticky traps were modeled after those described by Miles (1968). Dry ice was placed in a 1.9L cooler with 1 to 2 m of clear vinyl tubing (4.8 mm inside diameter and ∼10.1 mm outside diameter) inserted into the cooler drinking spout. At the sampling location, the tubing was secured to a natural stick about 0.5m to 1m long with duct tape and pointed into the burrow. At the terminal end of the tubing and stick, a 4cm strip of glue board (Trapper Max Mouse & Insect, Sheffield, UK) was attached creating a u-shape with the adhesive side on the interior surface (Supplemental Figure 1). The sticky trap was inserted into nests or burrows and the dry ice sublimation from the cooler caused CO2 release at the end of the tube, attracting hematophagous arthropods which, when approached, became immobilized on the adhesive part of the tape. Traps were left in the burrows for approximately 24 h, after which soft ticks were removed from the traps with forceps and placed in vials with 70% ethanol.

### 2.2 Camera trap data

Visual observations of animal activity using trail cameras around collection sites were made at GCSNA at Little Crevice and Mad Crow caves in 2105 and 2016 (Kim, 2017). At LANWR, Camera traps (Stealth Cam 2022 G42NG 32MP No-Glow Trail Camera) were set up between March and August 2023 facing directly at the burrows as well as nearby locations within 100m that directed towards openings well-traveled by animals based on tracks. Animal species in the pictures were identified morphologically to the lowest taxonomic unit.

### 2.3 Soft tick processing, DNA extraction, molecular barcoding

Soft ticks from GCSNA were processed as described earlier (Busselman et al., 2021). Soft ticks from LANWR were measured, sexed and identified using taxonomic keys (Guzmán-Cornejo et al., 2019; Pratt & Stojanovich, 1963) as *Ornithodoros turicata*. Soft ticks were surface sterilized in 10% bleach for 15 seconds followed by two rinses in DNAse free water for 15 seconds each. DNA was extracted from individual ticks randomly selected to represent all sizes and stages from all burrows. If there were more than 25 ticks collected, 25 were selected for extraction; all ticks were used if fewer than 25 were collected. For this, the MagMAX CORE Nucleic Acid Purification Kit (Applied Biosystems ThermoFisher Scientific, Waltham, Massachusetts, USA) was used following manufacturer instructions. Total nucleic acid was eluted into 50 µl of elution buffer. In addition to morphological identification, a subset of randomly selected soft ticks (but including specimens from all sites within GCSNA and LANWR) were subjected to a barcoding PCR targeting a ∼700bp fragment of the mitochondrial cytochrome c oxidase subunit I gene (COI) using primers LCO1490 (5’GGTCAACAAATCATAAAGATATTGG3’) and HCO2198 (5’TAAACTTCAGGGTGACCAAAAAATCA3’) (Folmer et al. 1994). For this, the FailSafe 2× premix buffer E (Biosearch Technologies, Petaluma, CA, USA) was used with 0.5 μL of enzyme mix, 1 μL of each primer (10 pmol) and 3 μL of DNA template in a 25 μL of final volume per reaction. Cycling conditions involved an initial denaturation at 94 °C for 3 min, 40 cycles at 94 °C for 30 s, 50 °C for 30 s and 72 °C for 30 s, and a final extension at 72°C for 8 min. PCR products were visualized in a 2% agarose gel, positive samples were purified with ExoSap-IT (USB Corporation, Cleveland, OH, USA) following the manufacture’s protocol, and bi-directionally sequenced by Eton Bioscience (San Diego, CA, USA). Sequences were quality-trimmed and assembled using Geneious Prime 2023.2.1 (Biomatters Inc. San Diego, CA, USA) and the consensus results were used as a query in the NCBI Basic Local Alignment Search Tool, BLASTn (Altschul et al., 1990)

### 2.4 Metabarcoding PCR and sequence analysis

PCR amplification of a ∼145bp fragment from the 12S rRNA mitochondrial gene using Kapa Taq DNA Polymerase (Roche Sequencing, Indianapolis, IN, USA) was performed. Both forward (5’CAAACTGGGATTAGATACC3’) and reverse (5’AGAACAGGCTCCTCTAG3’) primers (Humair et al., 2007; Kieran et al., 2017) had 10-base barcodes tags added. (Shokralla et al., 2014). Conditions for the metabarcoding PCR consisted of an initial denaturation at 95°C for 3 minutes, followed by 40 cycles of 30 seconds at 98°C, 60 seconds at 63°C, and 1 min at 72°C. PCR for every sample was performed in duplicate with individual barcoded primers for each replicate. If one of the two replicates for a sample amplified and the other did not, one more attempt was made to generate a second amplicon. Replicates were successfully generated for 162 samples (55 GCSNA and 107 LANWR). PCR products were submitted to the Texas A&M Institute for Genome Sciences and Society for library preparation and sequencing. Libraries were prepared using xGen^TM^ ssDNA & Low Input DNA Library Prep kit from Integrated DNA technologies (Coralville, IA, USA), and run on the NovaSeq6000 platform (Illumina, San Diego, CA, USA).

For analysis of sequencing reads, paired reads were matched using BBmerge (https://jgi.doe.gov/data-and-tools/software-tools/bbtools/). Merged reads were analyzed using Seekdeep (https://seekdeep.brown.edu/) (Hathaway et al., 2018). Barcodes and primers were trimmed out, sequences were filtered for a minimum length of 100 bp, and reads that had a quality score of less than 25 across 75% length were rejected. Filtered reads were binned on single base difference. Chimeras were marked and removed. When replicates were available for samples, only sequences that were present in both replicates were accepted. Resulting sequences were matched with host names using Basic Local Alignment Search Tool (BLAST; National Center for Biotechnology Information, US National Library of Medicine) using MegaBLAST parameters against the GenBank database (https://www.ncbi.nlm.nih.gov/genbank/) (Altschul et al., 1990). Sequences showing 99% or higher identity to database sequence were identified to species level. GenBank matches with identities between 94.79% - 99% were accepted when only one species of the genus occurs in the US according to the GBIF database (https://www.gbif.org/); this included Collared peccary, (*Pecari tajacu*), North American porcupine (*Erethizon dorsatum*), barking frog (*Craugastor augusti*), nine-banded armadillo (*Dasypus novemcinctus*), and Virginia opossum (*Didelphis virginiana*). In other instances where multiple species occur within Texas or the county of sampling, the lowest common taxonomic unit was assigned; this included Passeriformes – order level, Sciuridae – family level, *Crotalus* and *Neotoma* – genus level. Samples with fewer than 1000 total reads were removed from all analyses. The possibility of presence of a vertebrate host in the study area was taken into consideration. Hosts whose presence could not be established, given currently available information (GBIF database https://www.gbif.org/) were removed.. Since the target DNA is mitochondrial, chromosomal matches were also removed. After host identification, within each sample, vertebrate hosts with either fewer than 500 reads or fewer than 1% of the total reads for that sample, whichever higher, were removed. Many steps were taken to minimize contamination such using filter pipette tips and single use consumables, conducting all laboratory steps in disinfected biosafety cabinets and by including negative controls in all metabarcoding PCR runs, all of which were negative.

### 2.4 Diversity analysis and host network

To compare the diversity of hosts in each community identified through our metabarcoding approach, we calculated Simpson’s index of diversity for each cave or burrow within GCSNA and LANWR using the number of different hosts found in each site’s population of ticks:

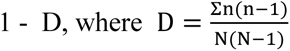

where n = number of individuals within each host species and N= total number of individuals identified across all hosts. A value closer to 1 indicates a more diverse host community and a value closer to 0 indicates a less diverse host community. This metric takes into consideration both the richness and evenness of species diversity. Diversity was then compared by averaging the Simpson’s index of diversity at each site within both locations, and a Mann-Whitney-U test was performed. A vector-host network to show the diversity of hosts associated with each collection site was created (Figure 1). Each site where *O. turicata* were found is listed along the left side, and all identified hosts are listed to the right. The width of the lines connecting site to host is proportional to the number of bloodmeal hosts identified at that site, and multiple lines from a site are used to identify all host species identified at that site. Statistical analyses and data visualization was performed in Program R ((R Core Team, 2023) (in R Studio) (“Posit,” 2023) with package bipartite (Dormann et al., 2008).

**Figure 1.**
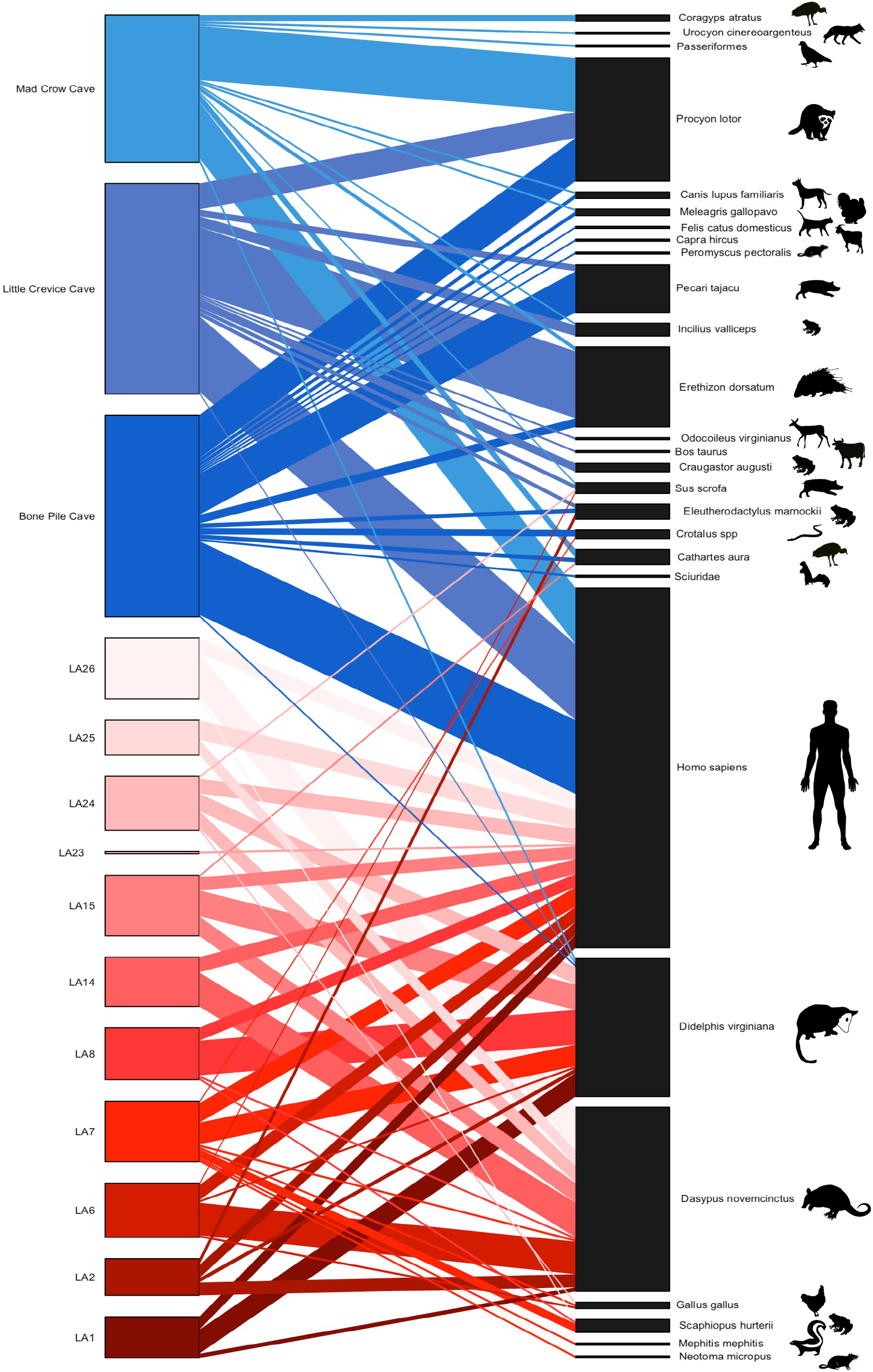
Graphic depiction of sites within Government Canyon State Natural Area (GCSNA) (left bars in blue) and Laguna Atascosa National Wildlife Refuge (LANWR) (left bars in red) and vertebrate hosts detected by bloodmeal metabarcoding (right column). The length of any given site’s bar is proportional to the number of hosts found and the lengths of the host bar reflects the number of times that hosts has been detected in bloodmeals in the study. The width of the networking lines indicate the number of times the host has been found at that location with the widest lines being the most frequent hosts for that location

## 3 RESULTS

### 3.1 Ornithodoros turicata collections

A total of a total of 381 soft ticks were collected from three caves within GCSNA, identified as *O. turicata*; 32 were female, 55 male, the rest were unidentifiable for sex, and 285 were nymphs, as previously reported in Busselman et al (2021). A total of 407 soft ticks were collected from 11 locations within LANWR from February to June, 2022. Specimens included 35 larvae, 235 nymphs, and 137 adults, which were morphologically identified as *O. turicata*. Molecular barcoding of 25 soft ticks – 8 from GCSNA (3 adults, 4 nymphs, 1 unknown) and 17 from LANWR (7 adults, 10 nymphs) – were all confirmed as *O. turicata*. Sequences binned into three *O. turicata* COI haplotypes that were 99.07-99.84% identical to each other (differences were 1, 5 and 6 bp). Two of these haplotypes (GenBank acc. num. XXXX and XXXX) were present in both locations, while the third one (GenBank acc. num. XXXX) was found only once in LANWR. These sequences matched GenBank *O. turicata* records with 99.07-99.8 % identity (GenBank acc. num. XXXX).

### 3.2 Metabarcoding analysis

From the 381 soft ticks collected at GCSNA, 168 (69 adults and 97 nymphs) were subjected to the metabarcoding PCR, products were visualized for 122 samples, 121 of which yielded at least 1000 final reads. Hosts were identified from 118 samples (47 adults, 70 nymphs, one unknown) at an average of 2.5 vertebrates identified as bloodmeal source per soft tick. Mammals were 14 of the 22 hosts, four were avian, one reptilian and three amphibian (Table1, Figure 1). After humans, the most frequent hosts were raccoon (*Procyon lotor*) followed by North American porcupine and collared peccary. The maximum number of hosts detected from one soft tick was six, (ST-47A) from Bone Pile cave: raccoon, collared peccary, turkey vulture (*Cathartes aura*), turkey (*Meleagris gallopavo*), human (*Homo sapiens*) and rattlesnake (*Crotalus* sp). (Supplemental spreadsheet GCSNA), while 5%, 19%, 58%, and 16% of the soft ticks had fed on four, three, two and one host, respectively.

Our previous work using Sanger sequencing (Busselman et al., 2021) for bloodmeal analysis in the same *O. turicata* specimens identified single-host DNA in 19 soft ticks out of 168 tested (11.3%), while here, host information was obtained from 72% of *O. turicata* tested by metabarcoding. This method revealed multiple host bloodmeals in 84% of the samples, adding 18 host species to the four different host species detected by Sanger sequencing (Table 2).

**Table 1:** Table of caves and hosts for Government Canyon State Natural Area (GCSNA), San Antonio, Texas, indicating the number of times each host was identified in *Ornithodoros turicata*. Visual confirmation is based on records from September 2015 to October 2016 (Kim, 2017).

FIGURES
| Vertebrates with visual confirmation | Overall | Bone Pile | Little Crevice | Mad Crow |
| --- | --- | --- | --- | --- |
| *Craugastor augusti*/Barking frog | 4 | 0 | 4 | 0 |
| *Coragyps atratus*/Black vulture | 3 | 0 | 0 | 3 |
| *Felis catus domesticus*/Cat | 1 | 1 | 0 | 0 |
| *Bos taurus*/Cattle | 1 | 0 | 1 | 0 |
| *Eleutherodactylus marnockii*/Cliff chirping frog | 4 | 2 | 2 | 0 |
| *Canis lupus familiaris*/Dog | 3 | 2 | 0 | 1 |
| *Sus scrofa*/Feral Hog | 3 | 0 | 2 | 1 |
| *Capra hircus*/Goat | 1 | 1 | 0 | 0 |
| *Urocyon cinereoargenteus*/Gray Fox | 1 | 0 | 0 | 1 |
| *Incilius valliceps*/Gulf coast toad | 6 | 0 | 5 | 1 |
| *Homo sapiens*/Human | 95 | 34 | 35 | 26 |
| *Erethizon dorsatum*/North American Porcupine | 36 | 4 | 30 | 2 |
| Passerifomes | 1 | 0 | 0 | 1 |
| *Pecari tajacu*/Collared peccary | 22 | 19 | 3 | 0 |
| *Procyon lotor*/Raccoon | 56 | 19 | 12 | 25 |
| *Crotalus* spp/Rattlesnake | 3 | 3 | 0 | 0 |
| Sciuridae | 1 | 1 | 0 | 0 |
| *Meleagris gallopavo*/Turkey | 3 | 2 | 0 | 1 |
| *Cathartes aura*/Turkey vulture | 6 | 2 | 0 | 4 |
| *Didelphis virginiana*/Virginia opossum | 4 | 1 | 1 | 2 |
| *Peromyscus pectoralis*/White-ankled mouse | 1 | 1 | 0 | 0 |
| *Odocoileus virginianus*/White-tailed deer | 1 | 0 | 1 | 0 |

**Table 2:**
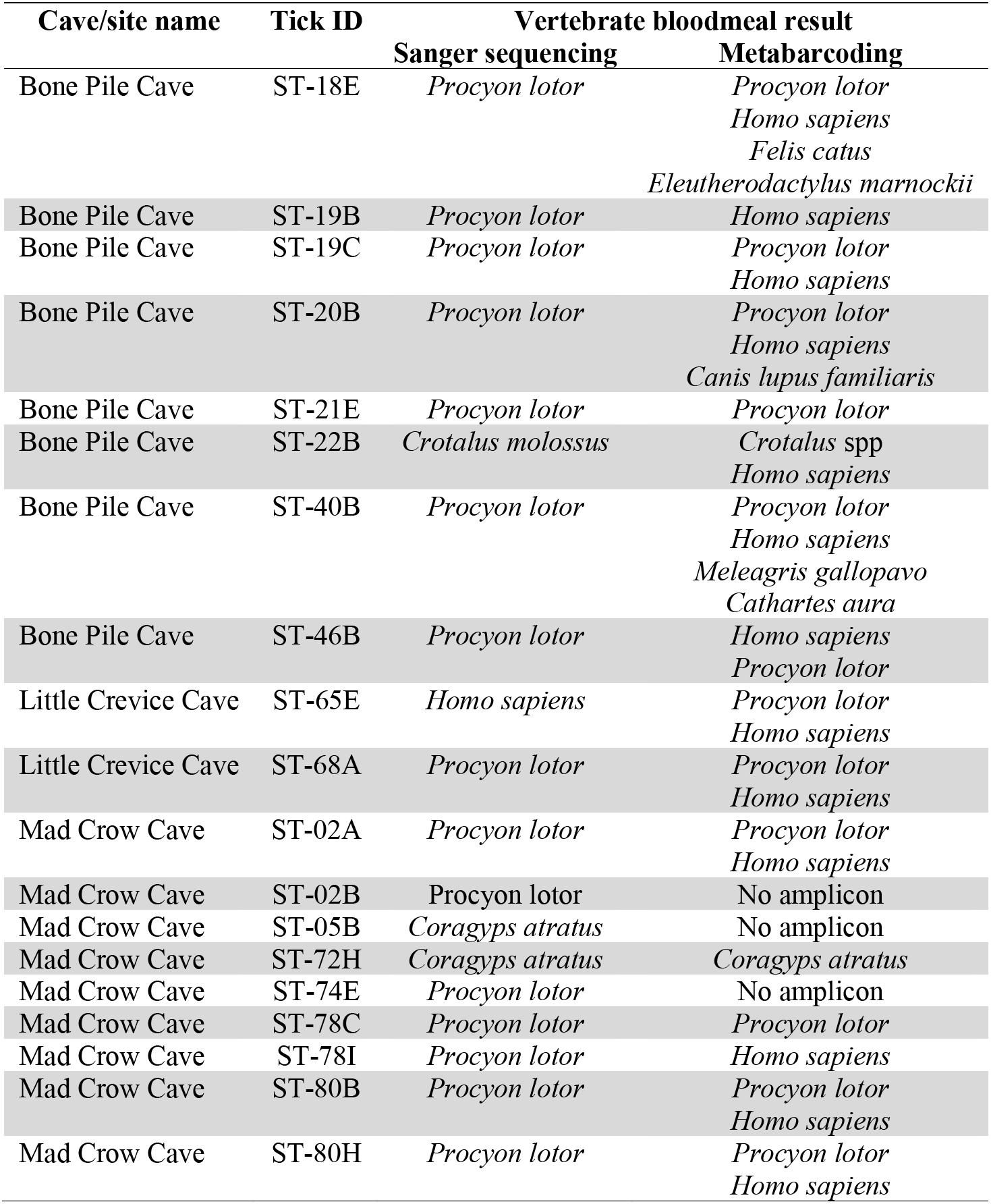
Comparison of results between Sanger sequencing (Busselman et al., 2021) and metabarcoding in *Ornithodoros turicata* collected at Government Canyon State Natural Area (GCSNA), San Antonio, Texas.

| Cave/site name | Tick ID | Vertebrate bloodmeal result |  |
| --- | --- | --- | --- |
|  |  | Sanger sequencing | Metabarcoding |
| Bone Pile Cave | ST-18E | *Procyon lotor* | *Procyon lotor*<br>*Homo sapiens*<br>*Felis catus*<br>*Eleutherodactylus marnockii* |
| Bone Pile Cave | ST-19B | *Procyon lotor* | *Homo sapiens* |
| Bone Pile Cave | ST-19C | *Procyon lotor* | *Procyon lotor*<br>*Homo sapiens* |
| Bone Pile Cave | ST-20B | *Procyon lotor* | *Procyon lotor*<br>*Homo sapiens*<br>*Canis lupus familiaris* |
| Bone Pile Cave | ST-21E | *Procyon lotor* | *Procyon lotor* |
| Bone Pile Cave | ST-22B | *Crotalus molossus* | *Crotalus* spp<br>*Homo sapiens* |
| Bone Pile Cave | ST-40B | *Procyon lotor* | *Procyon lotor*<br>*Homo sapiens*<br>*Meleagris gallopavo*<br>*Cathartes aura* |
| Bone Pile Cave | ST-46B | *Procyon lotor* | *Homo sapiens*<br>*Procyon lotor* |
| Little Crevice Cave | ST-65E | *Homo sapiens* | *Procyon lotor*<br>*Homo sapiens* |
| Little Crevice Cave | ST-68A | *Procyon lotor* | *Procyon lotor*<br>*Homo sapiens* |
| Mad Crow Cave | ST-02A | *Procyon lotor* | *Procyon lotor*<br>*Homo sapiens* |
| Mad Crow Cave | ST-02B | *Procyon lotor* | No amplicon |
| Mad Crow Cave | ST-05B | *Coragyps atratus* | No amplicon |
| Mad Crow Cave | ST-72H | *Coragyps atratus* | *Coragyps atratus* |
| Mad Crow Cave | ST-74E | *Procyon lotor* | No amplicon |
| Mad Crow Cave | ST-78C | *Procyon lotor* | *Procyon lotor* |
| Mad Crow Cave | ST-78I | *Procyon lotor* | *Homo sapiens* |
| Mad Crow Cave | ST-80B | *Procyon lotor* | *Procyon lotor*<br>*Homo sapiens* |
| Mad Crow Cave | ST-80H | *Procyon lotor* | *Procyon lotor*<br>*Homo sapiens* |

In the subset of 19 *O. turicata* that revealed hosts by Sanger sequencing, metabarcoding revealed that 11 had fed on multiple hosts, two of those feeding on four hosts each, and three soft ticks did not yield amplicons in the metabarcoding assay. While the Sanger sequencing approach yields only one host for all tested samples, only five of the same 19 soft ticks showed a single bloodmeal host by metabarcoding. Hosts identified by Sanger sequencing were also identified by metabarcoding in all but one instance (ST-78I).

Of the 407 *O. turicata* collected at LANWR, 226 (78 adults, 133 nymphs and 15 larvae) were tested via metabarcoding PCR and products were obtained from 159 samples. Information on bloodmeal hosts were obtained from 156 soft ticks (65 adults, 90 nymphs and one larva; 69% of the total ticks tested), revealing 11 host taxa across four vertebrate classes (Table 3). An average of 1.5 hosts per individual tick was identified. The nine-banded armadillo was the most frequently identified host, followed by human and Virginia opossum. A maximum of four different host species were identified in one soft tick from LANWR (LA-24R – nine-banded armadillo, human, chicken, Virginia opossum). Overall there was weak evidence of higher host diversity fed upon by *O. turicata* in GCSNA compared to LANWR (Simpson’s index of diversity values were 0.752 and 0.604, at GCSNA and LANWR respectively; p-value = 0.0769) (Muff et al., 2022).

**Table 3:** Trap site and hosts identified for Laguna Atascosa National Wildlife Refuge (LANWR), Los Fresnos, Texas, indicating the number of times each host was identified as a host. Visual confirmation is based on camera records from 2023 (see methods)

| Vertebrate host | Overall | LA14 | LA15 | LA1 | LA24 | LA25 | LA26 | LA2 | LA6 | LA7 | LA8 | Visual confirmation |
| --- | --- | --- | --- | --- | --- | --- | --- | --- | --- | --- | --- | --- |
| *Cathartes aura*/Turkey vulture | 1(0.4%) | 0 | 1(4%) | 0 | 0 | 0 | 0 | 0 | 0 | 0 | 0 | Y |
| *Crotalus* spp/Rattlesnake | 1(0.4%) | 0 | 0 | 0 | 0 | 0 | 0 | 0 | 0 | 1(4%) | 0 | N |
| *Dasypus novemcinctus*/Nine-banded armadillo | 85(36%) | 16(70%) | 9(32%) | 2(11%) | 8(32%) | 7(44%) | 20(71%) | 6(35%) | 15(60%) | 1(4%) | 1(4%) | Y |
| *Didelphis virginiana*/Virginia opossum | 60(26%) | 0 | 12(43%) | 11(58%) | 8(32%) | 0 | 0 | 2(12%) | 1(4%) | 10(36%) | 16(67%) | Y |
| *Eleutherodactylus marnockii*/Cliff chirping frog | 3(1%) | 0 | 0 | 0 | 0 | 0 | 0 | 3(18%) | 0 | 0 | 0 | N |
| *Gallus gallus*/Chicken | 3(1%) | 0 | 0 | 0 | 1(4%) | 0 | 0 | 0 | 1(4%) | 1(4%) | 0 | N |
| *Homo sapiens*/Human | 71(30%) | 7(30%) | 6(21%) | 6(32%) | 7(28%) | 9(56%) | 7(25%) | 6(35%) | 7(28%) | 9(32%) | 6(25%) | N |
| *Mephitis mephitis*/Striped skunk | 1(0.4%) | 0 | 0 | 0 | 0 | 0 | 0 | 0 | 0 | 1(4%) | 0 | Y |
| *Neotoma* spp/Woodrat | 1(0.4%) | 0 | 0 | 0 | 0 | 0 | 0 | 0 | 0 | 1(4%) | 0 | Y |
| *Scaphiopus hurterii*/Hurter's spadefoot toad | 6(3%) | 0 | 0 | 0 | 0 | 0 | 1(4%) | 0 | 0 | 4(14%) | 1(4%) | N |
| *Sus scrofa*/Feral hog | 2(0.9%) | 0 | 0 | 0 | 1(4%) | 0 | 0 | 0 | 1(4%) | 0 | 0 | Y |
| Total of vertebrate meals | 234 | 23 | 28 | 19 | 25 | 16 | 28 | 17 | 25 | 28 | 24 |  |

### 3.3 Integration with camera trap data

In total, 17 vertebrate species were identified both by metabarcoding and on camera, 16 were identified solely by metabarcoding and 23 were identified only on camera. At GCSNA, 11 of the vertebrates detected through metabarcoding were also observed on camera traps placed three years prior the tick sampling period for this study (Table 1, Supplemental Table 1) and 11 hosts were detected solely on metabarcoding. Ringtail (*Bassariscus astutus*), coyote (*Canis latrans*), Bobcat (*Lynx rufus*) and black-tailed jackrabbit (*Lepus californicus*) were observed on trail cameras but were not detected in bloodmeals. At GCSNA, counting both metabarcoding and camera trap vertebrates, 26 vertebrates were identified, with 22 of the 26 (85%) detected by metabarcoding alone. At LANWR, 30 vertebrates were identified by both methods combined - five by metabarcoding alone, six by both methods and 19 on camera alone so metabarcoding detected 11 of the 30 (37%) of all vertebrates combined. (Supplemental Table 1and 2)

## 4 DISCUSSION

This study is the first utilizing a metabarcoding approach to characterize the diversity of hosts fed upon by soft ticks, revealing a total of 27 different vertebrate species (22 in GCSNA, 11 in LANWR and 6 common to both locations) among bloodmeals of individual *O. turicata* sampled in Texas. Adult *O. turicata* have been documented to live over 7 years (Francıs, 1938), while those of *Argas brumpti* have been observed to live over 26 years (Shepherd, 2022). Such long-lived arthropods have opportunities to take multiple bloodmeals throughout their lifetime. Host DNA from bloodmeals remains viable in their abdomens for long periods and we have already documented the ability to detect bloodmeals of *O. turicata* over 3 years of age (Busselman et al., 2021), making them ideal arthropods to characterize vertebrate host communities by metabarcoding.

This study also allowed side-by-side comparisons between Sanger sequencing (see Busselman et al., 2021) and metabarcoding approaches in their efficiency to detect vertebrate hosts in bloodmeal analyses of *O. turicata*. In samples from GCSNA, metabarcoding showed an improvement in the rates of vertebrate detection in soft ticks from 11.3% to 72% of individuals tested and compared to Sanger sequencing. Overall, more than one host was detected in 168 of the of *O. turicata* analyzed (61% of samples for which bloodmeals were identified; 129 with two host, 31 with three hosts, seven with four hosts and one *O. turicata* with six hosts), while Sanger sequencing have revealed single-host bloodmeals only (Busselman et al., 2021; Kim et al., 2021; Palma et al., 2013), underscoring the advantages of the metabarcoding method to reveal cryptic vector-hosts interactions.

Although there was only weak evidence of differences in vertebrate community between GCSNA and LANWR based on Simpson’s index of diversity, only five overlapping hosts (turkey vulture, rattlesnake, Virginia opossum, cliff chirping frog (*Eleutherodactylus marnockii*) and wild pig) other than human hosts were detected in soft ticks in both locations (Figure1). This indicates differences in host communities between locations based on bloodmeal metabarcoding, which differed in the habitat where traps were deployed (caves and burrows), showing that host use varies at small spatial scales. Simpson’s index of diversity takes into consideration the richness and evenness of the community, and the average of each site’s diversity index within a location allowed statistical comparison between the two locations. Bloodmeal hosts identified from GCSNA showed differences in host feeding from cave to cave, with collared peccary being completely absent from Mad Crow cave and North American porcupine being more frequent in Little Crevice cave than the other two. Similarly, barking frog and most of the gulf coast toad (*Incilius valliceps*) bloodmeals are identified from Little Crevice cave (Table 1). In LANWR, while the most frequent bloodmeals were nine-banded armadillo or Virginia opossum, there are differences in the number of hosts between sites. The diversity of hosts covering four vertebrate classes and limited overlap in species detected between the two locations, confirms that *O. turicata* are opportunistic feeders with host choice likely varying with local availability of vertebrates.

At GCSNA, bloodmeal metabarcoding revealed host feeding on 22 vertebrate species out of the 26 species detected by camera traps and metabarcoding combined. At LANWR, only 11 species of hosts were detected in bloodmeals out of the 30 vertebrate species detected by camera traps and metabarcoding combined. Armadillos were reported as bloodmeal hosts in every LANWR burrow while camera traps only observed armadillos to be present in four of the burrows, and instead camera traps captured many additional vertebrates utilizing the burrows. This corroborates the recent camera trap observations of DeGregorio et al., (2022) that observed the importance of armadillos as ecosystem engineers providing shelter to diverse vertebrate communities. This study in Arkansas used camera traps to observe animals associated with armadillo burrows and found many vertebrate species entering and exiting the active nine-banded armadillo burrows; Virginia opossums co-inhabited the burrows with nine-banded armadillos. In our study, camera traps on LA14 documented concurrent use of the burrow by nine-banded armadillo and Virginia opossums (Supplemental Figure 2), while nine-banded armadillos and humans were the only vertebrates documented as *O. turicata* bloodmeal hosts from that burrow. A nine-banded armadillo was observed entering the LA8 burrow when soft tick traps were being set up at the same location. One year later, the camera trap on that burrow documented several additional vertebrates, including Virginia opossum; 67% of the tick bloodmeals from that burrow were from opossum with only 4% from armadillo. While much of the *O. turicata* host feeding appears opportunistic, we also observed frequent vertebrates associated with the burrows and fewer bloodmeals than we would expect. For example, eastern cottontails were observed at 6 of the LANWR camera trap locations, but no soft tick bloodmeals were observed from this species. Additionally, multiple species of rodents were frequently observed entering and exiting the burrows but only 1 out of 235 identified bloodmeals came from a rodent. Our use of camera trapping was not able to identify unique individuals of each species or estimate relative densities which precluded our ability to measure *O. turicata* host selection, or the over- and under- utilization of different vertebrates relative to abundance (Fikrig & Harrington, 2021). While *O. turicata* appears overall to be opportunistic, this study indicates that some vertebrates may over and under-utilized relative to availability, which could be pursued in future controlled studies.

*Ornithodoros turicata* is a vector of *Borrelia turicatae* in Texas and in Mexico, which causes tick-borne relapsing fever in humans and dogs (Beeson et al., 2023; Bissett et al., 2018). This study’s results identify the diverse community of potential reservoir hosts that could contribute to the enzootic maintenance of *B. turicatae* in nature. From the four mammal species detected as potential hosts of *B. turicatae* (Armstrong et al., 2018), two (raccoons and gray fox) were found in *O. turicata* bloodmeals. All these cases happened in caves where human and dog DNA was also detected. While acknowledging the potential for human bloodmeal results to originate from environmental DNA, the current study presents a potential mechanism of bridge transmission of *B. turicatae* from the enzootic cycle to humans and domestic animals. The evidence of bloodmeal derived from dogs in *O. turicata* sampled in caves, provide clues about the locations where *B. turicata* may be transmitted. Clinical cases of canine tick-borne relapsing fever have been reported in dogs in Texas (Piccione et al., 2016; Whitney et al., 2007). Human exposure to *O. turicata* bites leading to exposure to *B. turicatae* has been known to occur during brief periods of outdoor activity and while seeking refuge in caves (Davis et al., 2002; Dworkin et al., 2008; Forrester et al., 2015).

Evidence of soft tick feeding on feral hogs was not uncommon from these locations – 3 individual *O. turicata* at GCSNA and 2 at LANWR. *Ornithodoros turicata* have been identified as competent for the transmission of ASFV (Manzano-Román et al., 2012), and prior observations have documented the rare on-host observations of *O. turicata* on wild pigs (Sames & Teel, 2022). Texas is included in the region of the US with elevated risk for transmission of ASFV given the presence of competent vectors and wild pigs, domestic swine, and free-range common warthog (*Phacochoerus africanus*) documented in 4 counties (Mayer et al., 2020). The results of this study provides evidence of soft ticks and swine contact which could be occurring more frequently than previously known (Brown & Bevins, 2018; Herrera-Ibatá et al., 2017; Wormington et al., 2019), aided in part by molecular methodological advancements. In addition, the simple CO_2_-baited sticky trap deployed into the burrows in LANWR represents a technique that is not widely used for collecting softs ticks in endemic or non-endemic regions for ASFV (Jori et al., 2023). This technique can utilize dry ice, as in the current study, but also CO_2_ cylinders or sugar-yeast fermentation to increase the flexibility in diverse regions when resources are often limited.

In addition to *O. turicata* feeding on wild pig, we also observed 22 soft ticks from GCSNA to have fed on collared peccary. Although peccaries are in the new world pig family (Tayassuidae) they share the same suborder of Suina with old world pigs which are known to be highly susceptible to ASFV (e.g. *Sus scrofa* and *Phacochoerus* spp.). While peccaries are considered resistant to ASFV (Brown & Bevins, 2018), there are no experimental studies confirming this status. Consequently, the current observation of frequent feeding on peccaries by competent ASFV vectors warrants revisiting the ASFV vertebrate competence of new world pigs.

Human hosts were found to be the most frequently identified meal (95 out of 256) in GCSNA and the second most frequent (71 of 234) in LANWR. Humans are known to visit both the caves at GCSNA (camera trap evidence in Kim, 2017) and are known to recreate in the areas around burrows at LANWR. While it is plausible that some *O. turicata* human bloodmeal results are real, it is also possible that some may reflect environmental contamination. Human DNA is ubiquitous and is a component of all aspects of this study the placing of traps, tick collection, identification, extraction, and molecular steps. Despite surface sterilization of the soft ticks to remove exogenous DNA and measures taken during processing to avoid human contamination, the presence of human DNA contaminating soft tick samples cannot be ruled out. There is potential for some of these meals to be very old (Beck et al., 1986; Francıs, 1938; Kim et al., 2021). Future steps to parse sequences into human haplotypes to discern between different human encounters, contamination and meals would be useful.

We have collected unique information elucidating the feeding habits of soft ticks substantiating the diversity of hosts for *O. turicata* in natural areas. The detection of multiple meals within single ticks allows better deciphering of feeding over time, possibly across years, during which pathogens could remain viable. The metabarcoding protocol developed in this study is a valuable tool to detect vertebrate bloodmeal from soft ticks in areas currently affected by ASFV, identifying contact between susceptible hosts and local *Ornithodoros* vectors. Not only could bloodmeals be used to determine the transmission chain and inform control method for *B. turicatae* and ASFV, but also could inform the habits and presence of hosts.

## Supporting information

Supplemetal Spreadsheet GCSNA and LANWR

## ACKNOWLEDGEMENTS

This work was partially funded by a seed grant from the Coalition for Epi Response, Engagement, and Science (CERES), Texas A&M AgriLife Research, and Texas Ecolab. We also appreciate assistance sampling ticks and operating camera traps at Laguna Atascosa National Wildlife Refuge from Ester Carbajal, Danya Garza, Taylor Donaldson, Brian Rich, Odalis Sauceda, Javier Elizondo, Salvador Solis, Daniel, and Sarah Maestas.

## DATA ACCESSIBILITY AND BENEFIT-SHARING SECTION

### Data Accessibility Section

*O. turicata* barcoding sequences will be deposited into GenBank

Next generation sequencing data will be deposited in NCBI SRA or comparable public sequence read repository.

Voucher specimens of soft ticks identified as *O. turicata* from LANWR will be placed into Texas A&M University Insect Collection (https://entomology.tamu.edu/tamu-insect-collection/)

### Benefit-Sharing Section

Research at both locations in this study involved coordination with state level Texas Parks and Wildlife Department and Federal U.S. Fish and Wildlife Service and involvement of personnel from these agencies in our research. Local participants are acknowledged in the acknowledgements section.

## AUTHOR CONTRIBUTIONS

- Conceived the study: Sujata Balasubramanian, Rachel E. Busselman, Gabriel Hamer, Sarah Hamer, Pete D. Teel
- Performed research Sujata Balasubramanian, Rachel E. Busselman, Nadia Fernandez- Santos, Francisco C. Ferreira, Andrew Hillhouse, Nicholas Wolff
- Contributed new reagents or analytical tools Sarah Hamer, Gabriel Hamer, Nicholas Hathaway, Jeffrey A. Bailey
- Analysed data – Sujata Balasubramanian, Rachel E. Busselman, Nadia Fernandez-Santos, Andy Grunwald.
- Wrote the paper – Sujata Balasubramanian, Gabriel Hamer, Rachel Busselman, Francisco C. Ferreira
- Reviewed and approved submission – All authors

