## Supplemental Figures 1 & 2, Supplementa Tables 1 & 2 for "Bloodmeal metabarcoding of the argasid tick (*Ornithodoros turicata* Dugès) reveals extensive vector-host associations"

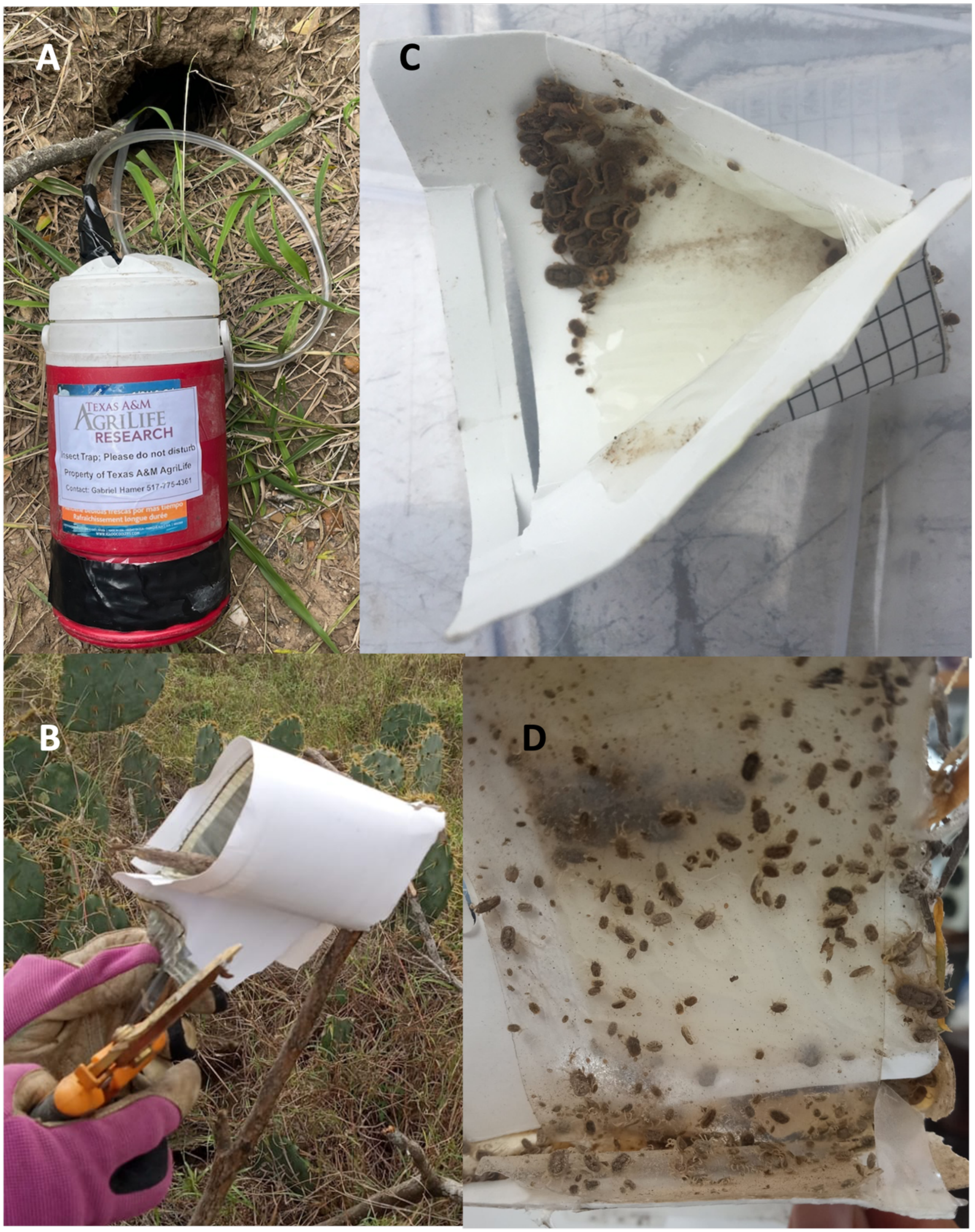


**Supplemental Figure 1.** $CO_2$-baited sticky traps used at Laguna Atascosa National Wildlife Refuge showing A) dry ice in the cooler with a tube inserted into burrow allowing sublimation of $CO_2$ to enter the burrow, B) sticky glue board wrapped around the end of the stick and tube so that sticky surface is on the inside, C) example of *O. turicata* stuck to the inside edge of the glue board, and D) example of *O. turicata* stuck to inside edge of sticky tape.

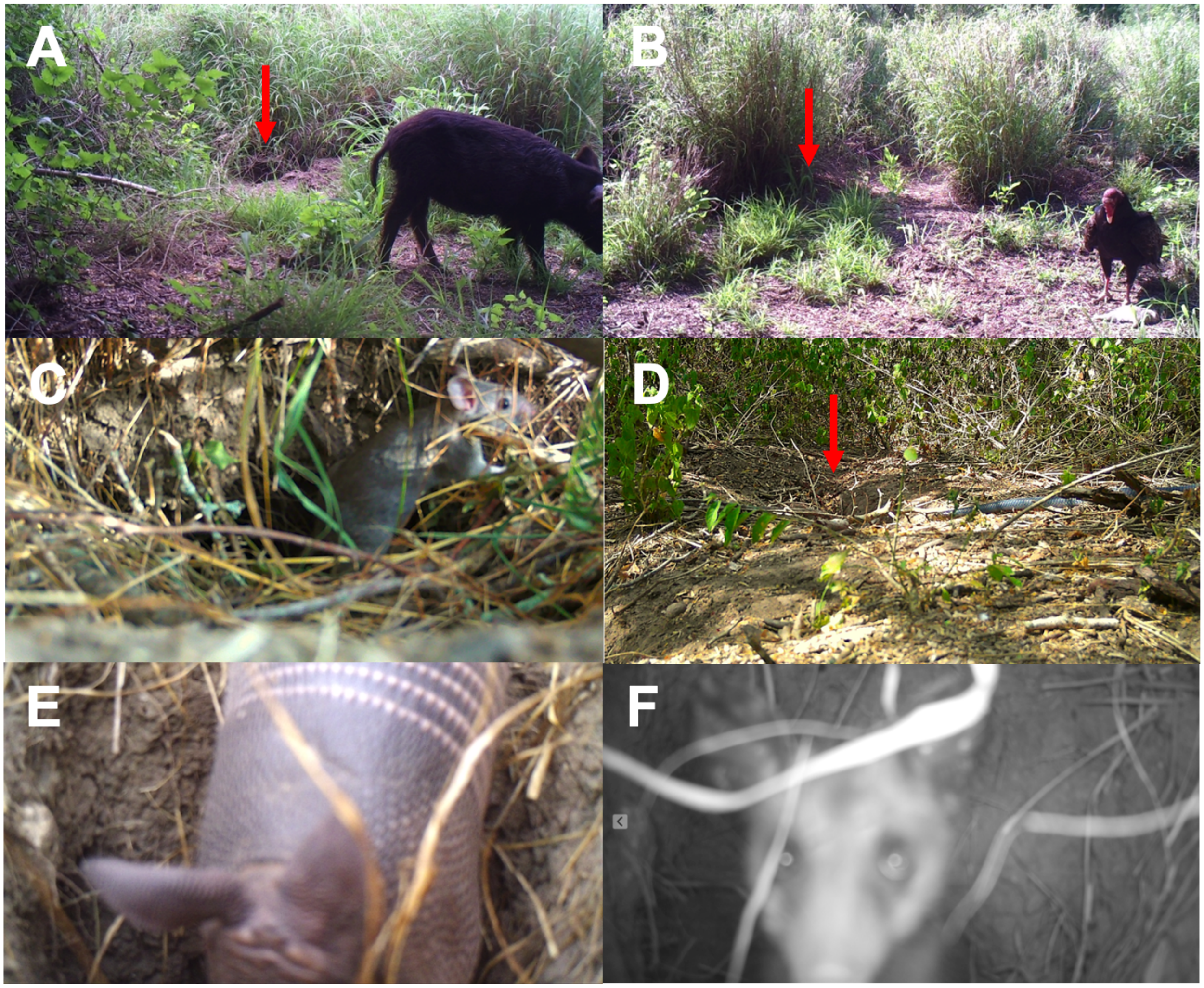


**Supplemental Figure 2.** Representative camera trap photos of A) wild pig (*Sus scrofa*) at burrow LA29, B) turkey vulture (*Cathartes aura*) at burrow LA29, C) Woodrat (*Neotoma* sp.) at burrow LA7, D) Indigo snake (*Drymarchon corais*) at burrow LA8, E) Nine-banded armadillo (*Dasypus novemcinctus*) at burrow LA14, and F) Virginia opossum (*Didelphis virginiana*) at burrow LA14. Photos E and F were taken one day apart at the same burrow. Red arrow pointing to the location of the burrows when the opening was not obvious

| | Metabarcoding only vertebrates | Metabarcoding And Camera vertebrates | Camera only vertebrates |
|---|---|---|---|
| **GCSNA** | *Bos taurus*/ Cattle<br>*Canis lupus familiaris*/Dog<br>*Capra hircus*/Goat<br>*Craugastor augusti*/Barking frog<br>*Eleutherodactylus marnockii*/Cliff chirping frog<br>*Felis catus*/Cat<br>*Incilius nebulifer*/Gulf coast toad<br>*Meleagris gallopavo*/Turkey<br>*Odocoileus virginianus*/White-tailed deer<br>*Pecari tajacu*/Collared peccary<br>*Peromyscus pectoralis*/White-ankled mouse | *Cathartes aura*/Turkey vulture<br>*Coragyps atratus*/Black vulture<br>*Crotalus* spp/Rattlesnake<br>*Didelphis virginiana*/Virginia opossum<br>*Erethizon dorsatum*/North American porcupine<br>*Homo sapiens*/Human<br>Passeriformes<br>*Procyon lotor*/Raccoon<br>Sciuridae<br>*Sus scrofa*/Feral hog<br>*Urocyon cinereoargenteus*/Grey Fox | *Bassariscus astutus*/Ring-tailed cat<br>*Canis latrans*/Coyote<br>*Lynx rufus*/Bobcat<br>*Lepus californicus*/Black-tailed jackrabbit |
| **LANWR** | *Crotalus spp*/Rattlesnake<br>*Eleutherodactylus marnockii*/Cliff chirping frog<br>*Gallus gallus*/Chicken<br>*Homo sapiens*/Human<br>*Scaphiopus hurterii*/Hurter's spadefoot toad | *Cathartes aura*/Turkey vulture<br>*Dasypus novemcinctus*/Nine-banded armadillo<br>*Didelphis virginiana*/Virginia opossum<br>*Mephitis mephitis*/Striped skunk<br>*Neotoma micropus*/Southern plains woodrat<br>*Sus scrofa*/Feral hog | *Leopardus pardalis*/Ocelot<br>*Lynx rufus/Bobcat*<br>*Canis latrans/Coyote*<br>Rodentia/Rodent<br>*Odocoileus virginianus/White-tailed deer*<br>*Boselaphus tragocamelus*/Nilgai<br>*Procyon lotor*/Racoon<br>*Sylvilagus floridanus*/Eastern cottontail<br>*Drymarchon corais*/Indigo snake<br>*Ictidomys parvidens*/Rio Grande ground squirrel<br>*Pecari tajacu*/Collared peccary<br>*Sceloporus olivaceus*/Texas spiny lizard<br>*Zenaida asiatica*/White-winged dove<br>*Toxostoma rufum*/Brown thrasher<br>*Caracara plancus*/Greater Roadrunner<br>*Ortalis vetula*/Plain Chachalaca<br>*Zenaida macroura*/Mourning dove<br>*Caracara plancus*/Crested caracara<br>*Thamnophis Proximus*/Western ribbon snake |

**Supplemental Table 1.** Hosts detected by different methods at Government Canyon State Natural Area (GCSNA) and Laguna Atascosa National Wildlife Refuge (LANWR). Visual recording of hosts at GCSNA was done from September 2015 to October 2016 (Kim, 2017) and LANWR from March to August, 2023.

| ID | | LA24 | LA8 | LA7 | LA14 | LA1 | LA2 | LA25 | | LA15 | | LA52 | LA49 | LA6 | | LA29 | |
|---|---|---|---|---|---|---|---|---|---|---|---|---|---|---|---|---|---|
| N | | 26.23234 | 26.23256 | 26.2323 | 26.2323 | 26.2332 | 26.2312 | 26.2323 | 26.1717 | 26.2323 | | 26.16 | 26.2329 | 26.2323 | 26.2163 | 26.1258. | |
| W | | 97.35201 | 97.35215 | 97.3521 | 97.352 | 97.3515 | 97.3516 | 97.3521 | -97.33 | -97.352 | | -97.321 | -97.352 | -97.352 | -97.352 | -97.211 | |
| Location type | | B | B | N | B | B | B | B | OS | B | *OS | N | B | B | OS | B | |
| Start date/ End date (2023) | | 08/02 to 08/05 | 7/17 to 7/25 | 7/17 to 7/25 | 7/17 to 7/25 | 7/17 to 8/2 | 7 /17 to 8/2 | 7/17 to 7/25 | 07/05 to 07/12 | 06/01 to 06/09 | 05/21 to 06/09 | 05/23 to 05/30 | 05/16 to 05/23 | 05/23 to 05/30 | 05/16 to 05/23 | 03/28 to 04/08 | |
| **Class** | **Animal/Days working** | 3 | 8 | 8 | 8 | 16 | 16 | 0 | 7 | 8 | 16 | 8 | 8 | 8 | 8 | 16 | Total |
| Birds | Brown thrasher *(Toxostoma rufum)* | | | | | | | | | | 1 | | | | 1 | | 2 |
| | Crested caracara *(Caracara plancus)* | | | | | | | | | | | | | | 1 | | 1 |
| | Greater roadrunner *(Geococcyx californianus)* | | | | | | | | | | 1 | | | | 1 | | 2 |
| | Mourning dove *(Zenaida macroura)* | | | | | | | | | | | | | | 1 | | 1 |
| | Plain chachalaca *(Ortalis vetula)* | | | | | | | | | | | | | | 1 | | 1 |
| | Turkey vulture *(Cathartes aura)* | | | | | | | | | | | | | | | 1 | 1 |
| | White-winged dove *(Zenaida asiatica)* | | | | | | | | | | 2 | | | | 1 | | 3 |
| Mammals | Bobcat (*Lynx rufus)* | | | | | | | | | | 1 | | | | 1 | 1 | 3 |
| | Collared peccary (*Tayassu tajacu* ) | | 1 | | | | | | | | | | | | | 3 | 4 |
| | Coyote *(Canis latrans)* | | | | | | | | | | 2 | | | | | | 2 |
| | Eastern cottontail (*Sylvilagus floridanus)* | | 1 | 1 | | | | | | | 1 | 1 | | | 1 | 1 | 6 |
| | Nilgai (*Boselaphus tragocamelus)* | | | | | | | | | | | | | | 2 | 2 | 4 |
| | Nine-banded armadillo (*Dasypus novemcinctus* ) | 1 | | 1 | 1 | | | 1 | | 1 | | | | | | 1 | 6 |
| | Ocelot *(Leopardus pardalis)* | | | | | | | | | | 1 | | | | | | 1 |
| | Raccoon *(Procyon lotor)* | | | | | | | | | | 1 | | | | | | 1 |
| | Rio Grande ground squirrel *(Ictidomys* | | | | | | | | | | 1 | 1 | | | | | 2 |
| | Rodent *(Rodentia* sp.) | 1 | 1 | 1 | | 1 | 1 | 1 | | | | 1 | 1 | 1 | | | 9 |
| | Striped skunk *(Mephitis mephitis)* | | | | | | 1 | | | | | | | | 1 | | 2 |
| | Virginia opossum *(Didelphis virginiana)* | | 1 | | 1 | 1 | | 1 | | | | | | | | 1 | 5 |
| | White-tailed deer *(Odocoileus virginianus)* | | | | | 1 | | | 1 | | 1 | | | | 2 | 3 | 8 |
| | Wild pig *(Sus scrofa)* | | | | | | | | | | | | | | | 1 | 1 |
| Reptiles | Indigo snake *(Drymarchon corais)* | | 1 | | | 2 | 1 | | | | | 1 | | | | | 5 |
| | Texas spiny lizard *(Sceloporus olivaceus)* | | 1 | | | | | | | | | | | | | | 1 |
| | Western ribbon snake (*Thamnophis proximus)* | | | | | | 1 | | | | | | | | | | 1 |
| | Total | 2 | 6 | 3 | 2 | 5 | 4 | 3 | 1 | 1 | 12 | 4 | 1 | 1 | 13 | 14 | 72 |

B- burrow, N-nest, OS-open site near burrows

* Location of camera trap with ocelot photo was removed given sensitivity of this federally endangered species

**Supplemental Table 2.** Camera trap observations at Laguna Atascosa National Wildlife Refuge, 2023. (Location of camera trap with ocelot photo was removed given sensitivity of this federally endangered species)
